# Regucalcin-containing extracellular vesicles suppress M2 macrophage polarization and attenuate tumor progression in vivo

**DOI:** 10.64898/2026.08.09.743746

**Authors:** Risako Okada, Kana Tominaga, Takeshi Yamamoto, Masayoshi Yamaguchi, Naoomi Tominaga

## Abstract

Regucalcin (RGN) plays diverse roles in cell biology, highlighting its importance in both physiological and pathological conditions. Prostate cancer patients with higher RGN expression exhibited significantly longer disease-free survival. Although RGN is a cell signaling suppressor, the molecular mechanisms underlying tumor suppression by RGN in the tumor microenvironment through cell–cell communication remain unclear. PC3 prostate cancer cell lines stably expressing RGN or a control vector were generated for this study. Extracellular vesicles (EVs) were isolated from these cell lines using differential ultracentrifugation. The murine macrophage cell line J744α1 was treated with isolated EVs, and effects on M2 polarization were evaluated using qRT-PCR and western blot analysis. To assess the potential anti-tumor effects of EVs, PC3 parental cells were subcutaneously implanted at two sites per mouse, followed by intratumoral injection of the respective EVs. Tumor volume was monitored. Harvested fresh frozen tumor tissues underwent immunofluorescence staining for CD206, an M2 macrophage marker. RGN was detected in EVs from RGN-expressing cells, and treatment with these RGN-containing EVs was associated with reduced tumor growth and reduced M2 macrophage polarization *in vitro* and *in vivo*. Furthermore, recombinant RGN protein reduced the levels of p-AKT1 and p-ERK1/2. Moreover, the suppression of M2 macrophage polarization by RGN-containing EVs was accompanied by decreased p-AKT1 and p-ERK1/2 *in vitro*. This study describes an EV-associated mechanism that may contribute to the regulation of macrophage polarization and indicates that RGN-containing EVs merit further evaluation as a candidate approach for cancer treatment. Causal validation, such as macrophage depletion or CD206 knockdown, and evaluation in additional models remain to be addressed in future studies.

## Introduction

Prostate cancer, the second most common cancer among men, leads to 7.3 cancer-related deaths per 100,000 men globally ^1^. Prostate cancer frequently metastasizes to bones, particularly the spine, pelvis, and ribs. Bone metastasis is the most common site of spread, affecting approximately 80% of patients during the disease course ^2^. Visceral metastasis, involving the lungs, liver, and adrenal glands, is less frequent but more aggressive and observed in 10%–20% of patients ^3^. Recent advances in understanding the molecular mechanisms of prostate cancer have driven the development of targeted therapies and improved diagnostic tools. Early detection and treatment are essential for enhancing patient outcomes ^4^. The prostate cancer tumor microenvironment (TME) is a complex and dynamic network that significantly impacts tumor progression and therapeutic resistance. It comprises various cellular components, including immune, stromal, and endothelial cells and the extracellular matrix, which interact with cancer cells to influence disease outcomes ^5^. Immune cells within the TME contribute to immunosuppressive conditions that promote tumor growth and metastasis. For example, tumor-associated macrophages (TAMs) and myeloid-derived suppressor cells (MDSCs) can suppress antitumor immune responses, facilitating cancer progression ^6^. Targeting the TME holds potential for improving patient outcomes by disrupting the supportive environment necessary for cancer cell survival and growth.

Regucalcin (RGN) plays a crucial role in maintaining intracellular calcium homeostasis and modulating diverse cellular processes in the body. Widely expressed across vertebrate and invertebrate species, RGN is recognized for its multifunctional properties as a cell signaling suppressor ^7^. The pleiotropic functions of RGN in cellular biology underscore its significance in both normal physiological processes and pathological conditions ^8^. Extracellular RGN, detected in human serum, has been shown to attenuate human cancer cell growth and bone metastasis ^9^. In prostate cancer, patients exhibiting elevated RGN expression levels demonstrated significantly prolonged recurrence-free and overall survival ^9^. Co-culturing RGN-expressing PC3 cells with pre-osteoblasts inhibits mineralization ^9^. Furthermore, co-culture with RAW264.7 cells suppressed osteoclast differentiation. Contemporary research indicates that inflammatory macrophages can mediate the destruction of prostate cancer cells via the NF-κB, STAT3, or RGN signaling pathways. These findings suggest that RGN expression in cancer cells mitigates cancer malignancy by suppressing macrophage polarization within the TME. Macrophages within the TME may exert a regulatory influence on the progression and metastasis of prostate cancer cells, potentially interacting with cancer cells to impede disease progression. However, the intricate interplay between macrophages and RGN-expressing prostate cancer cells remains largely uncharacterized.

Extracellular vesicles (EVs) are nanometer-sized lipid bilayer vesicles that encapsulate functional molecules such as mRNAs, miRNAs, DNA, and proteins. EVs mediate crucial cell-cell communication during various biological processes. For instance, miRNAs within breast cancer cell-derived EVs disrupt the integrity of the blood-brain barrier, thereby promoting brain metastasis ^10^. Pharmacological blockade of cancer cell-derived EVs in tumor-bearing mice has been shown to attenuate metastatic dissemination ^11^. Tumor-derived EVs modulate macrophage polarization, influencing tumorigenesis ^12^. EVs play a pivotal role in cancer progression by facilitating intercellular communication through the transfer of bioactive molecules ^13–15^. These molecules enclosed within EVs function as both tumor suppressors and oncogenes, thereby modulating gene expression in recipient cells ^11,15^. In recent years, EVs have garnered significant attention due to their potential application as cancer therapeutics ^13^. This study investigated the functional role of EVs derived from RGN-overexpressing prostate cancer cells in cancer progression. We detected RGN in EVs, which was accompanied by changes in their molecular cargo. Moreover, we found that RGN-containing EVs attenuate M2 macrophage polarization in vitro and that their administration was associated with reduced tumor growth *in vivo*.

## Material and Methods

### Public Data Analysis

Gene expression data and corresponding clinicopathological information of prostate cancer patients were acquired from Gene Expression Omnibus (GEO) database (https://www.ncbi.nlm.nih.gov/geo/). GSE94767 ^16^ and GSE29079 ^17,18^ provided RGN mRNA expression data from prostatic samples of patients with prostate cancer or healthy controls. GSE46602 ^19^ provided RGN mRNA expression data in benign and malignant prostate tissues. GSE6919 ^20,21^ dataset contains 152 human samples including primary or metastatic prostate cancer tissues, prostate tissues adjacent to the tumor, and organ donor prostate tissues. GSE40272 ^22^ dataset was used to compare the survival curves for disease-free survival generated by the Kaplan-Meier method with R package, ‘survival’ and ‘survminer’.

### Cell lines and cell culture

The human prostate cancer cell line, PC-3 (RCB2145), was provided by the RIKEN BRC through the National Bio-Resource Project of the MEXT/AMED, Japan on 2022∼2024. The cell lines authentication was performed by the RIKEN BRC. PC-3 was cultured in RPMI1640 medium (Nacalai Tesque, Japan, No.30264-85) supplemented with 10% heat-inactivated FBS (Corning, No.35-079-CV) and antibiotic–antimycotic agents (Nacalai Tesque, Japan, No.09366-44) at 37°C in 5% CO_2_. Established PC3-Emp and PC3-RGN cell lines were maintained in RPMI-1640 medium supplemented with 10% heat-inactivated FBS, 1% antibiotic–antimycotic agents (Nacalai Tesque, Japan, No. 09367-34), and 50 μg/mL G418 (Nacalai Tesque, Japan, No.09380-86). J744α1, the mouse macrophage cell line, was cultured in DMEM (Nacalai Tesque, Japan, No.16971-55) supplemented with 10% heat-inactivated FBS and antibiotic–antimycotic agents at 37°C in 5% CO_2_.

### Establishment of Cell lines

RGN expression and Empty vectors were kindly provided by Dr. Murata, Meijo University. PC-3 cells were seeded at a density of 5.0×10^5^ cells/well in 6-well plates and incubated for 24-hrs. PC-3 cells were transfected with plasmid DNA (2.0 μg) using calcium dichloride (120 mmol/mL, Fujifilm-Wako, Japan, No.038-24985) in HEPES buffer (Nacalai Tesque, No.17557-94) with media change after 4-hrs of incubation, and change to the culture medium. After 2 days of transfection, 500 μg/mL G418 was added to the culture medium to select transfected cells. The proliferating cells were reseeded in a 96-well plate using the limiting dilution method. Single cells were cultured and maintained in a culture medium containing 50 μg/mL G418.

### EVs isolation

Cells were seeded onto 15 cm dishes at 70-80% confluency and cultured overnight in a 5% CO_2_ incubator. The medium was then replaced with Advanced RPMI-1640 medium (Gibco, Massachusetts, No.12633020) supplemented with 2 mM L-glutamine (Nacalai Tesque, Japan, No.16948-04) and 1% antibiotic-antimycotic solution (Nacalai Tesque, Japan, No. 09367-34) after two washes with 10 mL Dulbecco’s Phosphate Buffered Saline (-) (Nacalai Tesque, Japan, No. 14249-24).

Following a 48-hour incubation period, the culture supernatant was collected by centrifugation at 10,000 xg for 10 min at 4°C. To ensure complete removal of cellular debris, the supernatant was subsequently filtered through a 0.22 μm PVDF filter (Millipore Merck, Massachusetts, No.17461-05). The resulting filtered supernatants were used for the isolation of EVs. EVs were isolated by ultracentrifugation at 110,000 xg for 70 min at 4°C using a SW41Ti swinging-bucket rotor (Beckman Coulter). The pellets were then washed with PBS(-) by repeated ultracentrifugation at 110,000 xg for 70 min at 4°C and resuspended in PBS(-). The isolated EVs were stored at 4°C until further experimentation.

### Dynamic Light Scattering (DLS)

The concentrated EVs samples were measured using a dynamic light scattering (DLS) instrument (ELSZneo, Otsuka Electronics, Co., Ltd., Osaka, Japan). The instrument was equipped with a 40 mW and 638 nm semiconductor laser diode. The laser was detected using an avalanche photodiode (APD) module. The scattering angle was 165◦, at a constant temperature of 25 °C. The optimal light-scattering intensity is between 84 cps and 114 cps (photon count rate per second). This system is equipped with an attenuation filter that automatically regulates the incident light if the scattering intensity is above the range. Each measurement consisted of 30 runs. The measurements were performed three times.

### Transmission electron microscopy (TEM)

The morphology of the EVs was confirmed using transmission electron microscopy ^23^. A formvar/carbon-coated copper grid (Nisshin EM, Tokyo, Japan) was hydrophilized with JFC-1600 Auto Fine Coater (JEOL, Tokyo, Japan). 3µL of purified EVs in PBS were placed on a hydrophilized grid and allowed to absorb for 3 min. We washed them using 500 µL double-distilled H_2_O successive four drops, then negatively stained using 30 µL of 2.0% uranyl acetate successive four drops. The grid was air-dried after absorbing 2.0% uranyl acetate on the grid using filter paper. We imaged the grid with Tecnai G2 Spirit BioTWIN electron microscope (FEI, OR, USA) operating 120 kV equipped with an EMSIS Phurona CMOS Camera. After exporting the raw data images to TIFF format, we used Fiji (ImageJ 1.53t) for data analysis.

### Western blotting

Proteins were isolated from cells using M-PER (Thermo Scientific, MA, USA) separated in Mini-PROTEAN TGX Gel (4–12%, Bio-Rad) and electro transferred onto a PVDF membrane (Bio-Rad, No.1704274). After blocking in Block Ace (KAC, Japan, No.UKB80), the membranes were incubated for 1-hr at room temperature with primary antibodies, which included anti-Regucalcin (Rabbit polyclonal, HPA029103-100ul, 1:500, Sigma), anti-GAPDH (Mouse monoclonal, clone 6C5, CB1001, 1:5000, Millipore), anti-β-actin (Mouse monoclonal, clone 2F1-1, No.643802, 1:2000, Biolegend), anti-CD63 (purified mouse anti-human CD63, H5C6, 1:200, BD), anti-Cytochrome C (Rabbit polyclonal, 4272S, 1;1000, CST), anti-Stat1 (D1K9Y) (Rabbit monoclonal, No.14994, 1:1000, CST), anti-Phospho-Stat1(Y701) (Rabbit monoclonal, No.7649, 1:1000, CST), anti-Phospho-Stat1(S727) (Rabbit monoclonal, No.8826, 1:1000, CST), anti-AKT1 (Rabbit polyclonal, No.10176-2-AP, 1:5000, Proteintech), anti-Phospho-AKT1 (Ser473) (Rabbit polyclonal, No.28731-1-AP, 1:3000, Proteintech), anti-ERK1/2 (Rabbit polyclonal, No.11257-1-AP, 1:5000, Proteintech), anti-Phospho-ERK1/2 (Thr202/Tyr204) (Rabbit polyclonal, No.28733-1-AP, 1:5000, Proteintech), anti-NFκB (Rabbit polyclonal, clone.Poly6226, No.622602, 1:5000, Biolegend), anti-Phospho-NFκB (Ser536) (93H1) (Rabbit monoclonal, No.3033, 1:1000, CST). Secondary antibodies (HRP-linked anti-mouse IgG, NA931V or HRP-linked anti-rabbit IgG, NA934V, GE Healthcare) were used at a dilution of 1:10000. The membrane was then exposed to ImmunoStar LD (Fujifilm-Wako, Japan, No.292-69903).

### Cell proliferation assay

PC-3 (PC3-Parental), Emp3 (PC3-Emp), and RGN3 (PC3-RGN) cells were seeded at a density of 1×10^4^ cells per well in a 96-well plate. The cell proliferation was assessed cell number using a hemocytometer. The cell proliferation was measured on days 0, 1, 2, 3, and 5.

### Mouse studies

A cell suspension containing 2×10^5^ PC-3 prostate cancer cells in a volume of 100 μL of Matrigel (Corning, No.354234) was injected into the subcutaneous dorsal side of anaesthetized 7-weeks-old BALB/c Slc-nu/nu male mice. Cells were implanted at two dorsal sites per mouse, and the implantation site was used as the experimental unit for analysis.

Two independent in vivo experiments were performed. In the tumor formation experiment (Fig. 2), parental PC3, PC3-Empty, or PC3-RGN cells were implanted using three mice per group, giving six implantation sites per group, and tumors were followed for 35 days. In the EV treatment experiment (Fig. 6), parental PC3 cells were implanted in all animals, and mice were assigned to four treatment groups (no treatment, parental PC3 EVs, PC3-Empty EVs, or PC3-RGN EVs) using three mice per group, giving six implantation sites per group. EVs were administered by intratumoral injection at 5 μg per site on days 7, 13, 16, and 19 after implantation, for a total of four injections, and tumors were followed for 22 days.

Tumor development was monitored by 3-days intervals. Tumor dimensions were measured with calipers. After euthanasia, the tissues were rapidly excised and embedded in Optimal Cutting Temperature (OCT) compound, followed by snap-freezing in liquid nitrogen and processed for histological analysis. Tumor volume was calculated using the formula: V=0.5・(longest diameter)・(shortest diameter)^2. The tumors were halved, and the cut surfaces were embedded in a face-down position. All tissues were stored at -80 °C until used for staining and imaging experiments. One PC3-RGN implantation site failed to form a palpable tumor and was included in the analysis with a volume of 0 mm³; no animals or data points were otherwise excluded. All animal experiments followed a protocol approved by the Yamaguchi University Animal Care and Use Committee (No.80-021).

### Immunofluorescence assay

The tumors in OCT compound were sectioned to a thickness of 10 μm, then the tissue sections were mounted onto glass slides (Matsunaim Glass Ind., Japan, No.SCRE-15). The sectioned tissue was washed in PBS to remove OCT compound, then fixed with 4% paraformaldehyde in PBS for 10 min at room temperature and treated with PBS containing 0.1% Triton X-100 for 10 min after washing with PBS (-) to permeabilize the cells. After fixation, the cells were incubated with 3% BSA in PBS for 1-hr to block the nonspecific binding of antibodies. Subsequently, the tumor samples were incubated with rat monoclonal APC-conjugated antibody against CD206 (MR6F3, No.17-2061-82, 1:100, ThermoFisher Scientific), rabbit polyclonal antibody against Regucalcin (HPA029103, 1:500, Sigma-Aldrich), or mouse monoclonal antibody against Vimentin (No.677801, 1:500, Biolegend) suspend in Can Get Signal immunostain Immunoreaction Enhancer Solution A (NKB-501, TOYOBO, Japan) at 4 °C for overnight. After washing with PBS containing 0.1% Triton X-100, anti-RGN antibody treated samples were incubated with Alexa Fluor 488-conjugated anti-rabbit IgG (No.A11034, 1:500, Invitrogen) suspend in Can Get Signal immunostain Immunoreaction Enhancer Solution B (NKB-601, TOYOBO, Japan) for 1-hr at room temperature. The stained cells were then washed in PBS containing 0.1% Triton X-100 and were mounted in ProLong Glass Antifade Mountant (P36984, ThermoFisher Scientific) for observation under a confocal microscope (BZ-X810, Keyence, Japan). For nuclear staining, 4’,6-Diamidino-2-phenylindole (DAPI, D523, Dojindo) was used.

### Protein transfection

J744α1 cells were cultured in 6-well plates at a density of 6.0 × 10^^5^ cells per well. The following day, 4 ng/μL of recombinant Human-RGN (No. pro-915a, PROSPEC) was transfected using ProteoCarry (No. FDV-0015, Funakoshi) after a PBS (-) wash. The cells were incubated for 1 hour at 37°C in 5% CO_2_. After 1 hour, the transfection mixture was removed, the cells were washed three times with PBS (-), and then the culture medium was added. mRNA was extracted on day 2 after transfection.

### Quantitative real-time PCR

J744α1 cells were cultured in 6-well plates at a density of 6.0×10^5^ cells per well. The following day, the cells were treated with EVs (1 µg/mL) obtained from untransfected PC3 (PC3-Parental), control empty vector-transfected PC3 (PC3-Emp), and RGN-overexpressing PC3 cells (PC3-RGN), and incubated for 24 hours. Untreated PC3 cells were used as a control (N.C.). Total RNA was extracted using the ISOSPIN Cell & Tissue RNA kit (Nippon Gene, Tokyo, Japan) and reverse transcribed into cDNA using the ReverTra Ace qPCR RT Master Mix (TOYOBO, Osaka, Japan). Quantitative PCR (qPCR) was performed in triplicate using the PowerTrack SYBR Green Master Mix (Invitrogen) on a QuantStudio 3 real-time PCR system (Applied Biosystems, USA). GAPDH was used as the reference gene for relative quantification. The primers used in this study are listed in the Table 1. Product specificity was confirmed by melting curve analysis at the end of each run, and a single peak was obtained for each primer pair. Gene expression levels were quantified using the standard curve method.

**Table 1.**
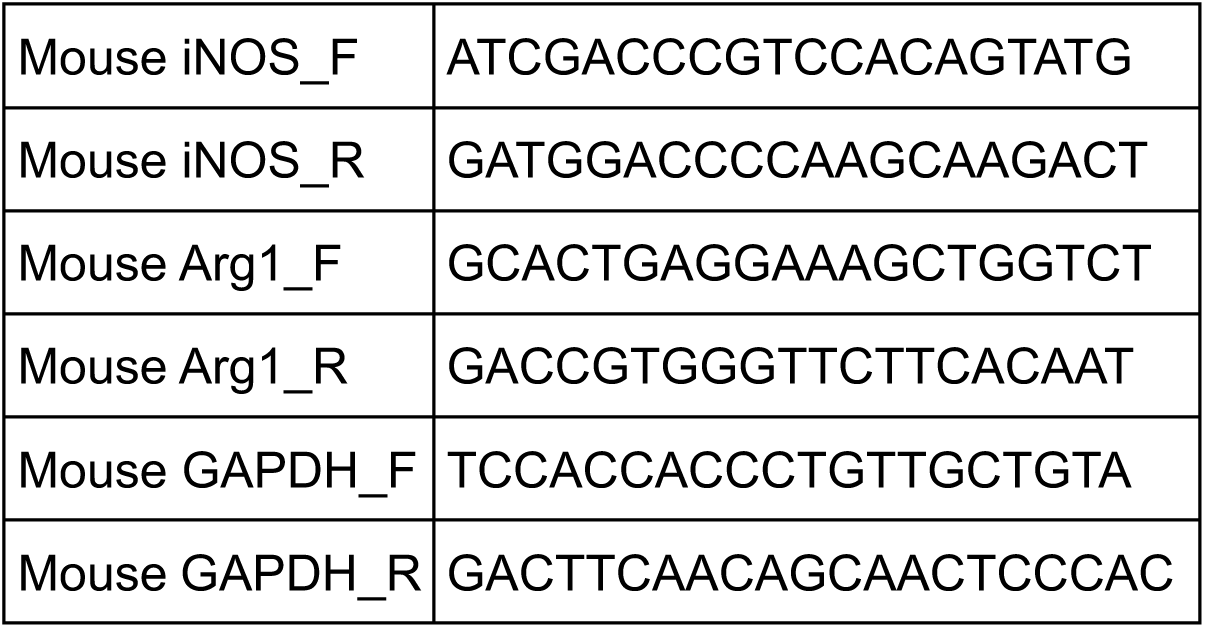
Primers for qPCR.

### Quantification and statistical analysis

We conducted statistical analyses and visualized the data utilizing R software (version 4.2.1). Normality of the data was assessed using the Shapiro–Wilk test prior to parametric testing. To compare differences between two groups, we employed Student’s t-test. For comparisons among three or more groups, one-way ANOVA followed by the Tukey–Kramer post-hoc test was used. P values below 0.05 were considered statistically significant. Graphical representations display the mean ± standard deviation (S.D.).

## Results

### Reduced RGN in malignant tissues correlates with advanced tumor stages and decreased survival

To elucidate the role of RGN in tumorigenesis, we analyzed its expression profiles across multiple independent microarray datasets. RGN mRNA levels were significantly downregulated in tumor tissues compared to normal tissues in all three datasets (Fig. 1a, b, and c). Specifically, in the GSE94767 dataset, RGN expression in tumor samples was substantially reduced (p = 6.7 × 10^−6^), demonstrating a clear differential expression pattern. Similarly, the GSE29079 dataset revealed a marked decrease in RGN expression within tumor samples (p = 7.4 × 10^−7^). The GSE46602 dataset, which included benign and tumors, exhibited significant downregulation of RGN in tumor samples (p = 0.0012). These findings suggest an association between RGN downregulation and malignant transformation. Further analysis of the GSE6919 dataset (Fig.1d) revealed the dynamic expression of RGN across various stages of tumor progression: normal tissue, adjacent normal tissue, primary tumor, and metastatic tumor. Kruskal-Wallis test with Dunn’s post-hoc correction revealed a progressive decline in RGN expression, with the lowest levels detected in the metastatic tissues. This progressive downregulation suggests that lower RGN expression is associated with more advanced disease stages in these datasets. Log-rank test survival analysis using the GSE40272 dataset (Fig.1e) revealed a significant inverse correlation between RGN expression and disease-free survival (p = 0.017). Patients with lower RGN expression exhibited significantly shorter survival times than those with higher expression. The number-at-risk table showed a greater rate of attrition in the low RGN expression cohort. These data indicate that reduced RGN expression is associated with tumor progression and shorter disease-free survival in these patient cohorts.

**Figure 1.**
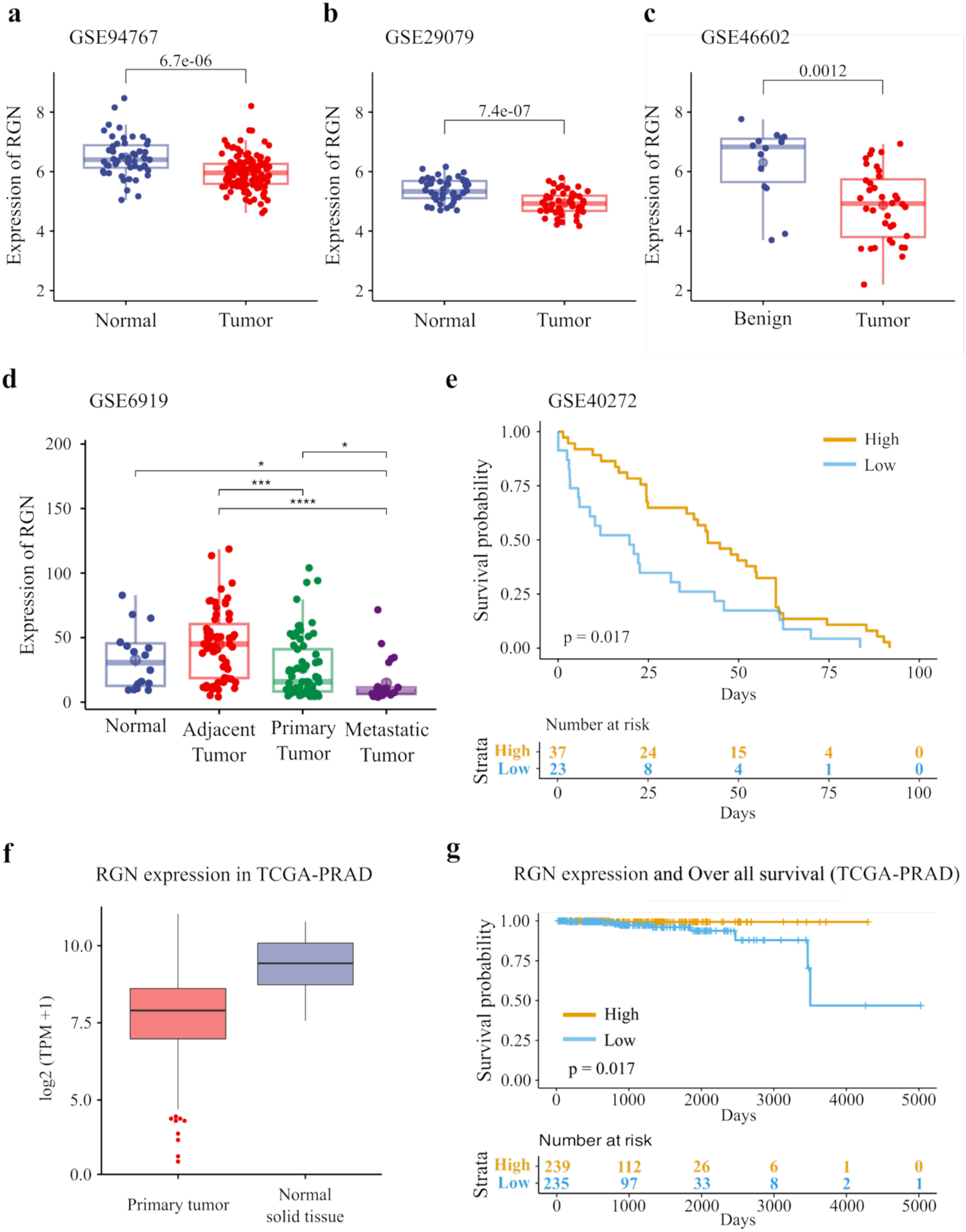
RGN expression patterns and clinical significance. **a-c.** Box plots comparing RGN expression between normal and tumor tissues (GSE94767, GSE29079) or benign and tumor tissues (GSE46602). Mann–Whitney U test P-values are shown. **d.** Box plot of RGN expression across normal, adjacent, primary, and metastatic tumor tissues (GSE6919). Kruskal–Wallis test with Dunn’s post-hoc test; \**P* < 0.05, \*\**P* < 0.01, \*\*\**P* < 0.001. e. Kaplan-Meier survival curves for high and low RGN expression (GSE40272). Log-rank test P-values and patient risk table are shown. f. Box plot comparing RGN expression between primary tumor and normal solid tissue in the TCGA-PRAD cohort. Expression is shown as log2(TPM+1). P-value is shown. g. Kaplan-Meier curves for overall survival in the TCGA-PRAD cohort, stratified by RGN expression into high (n = 239) and low (n = 235) groups. Log-rank test P-value and patient risk table are shown.

**Figure 2.**
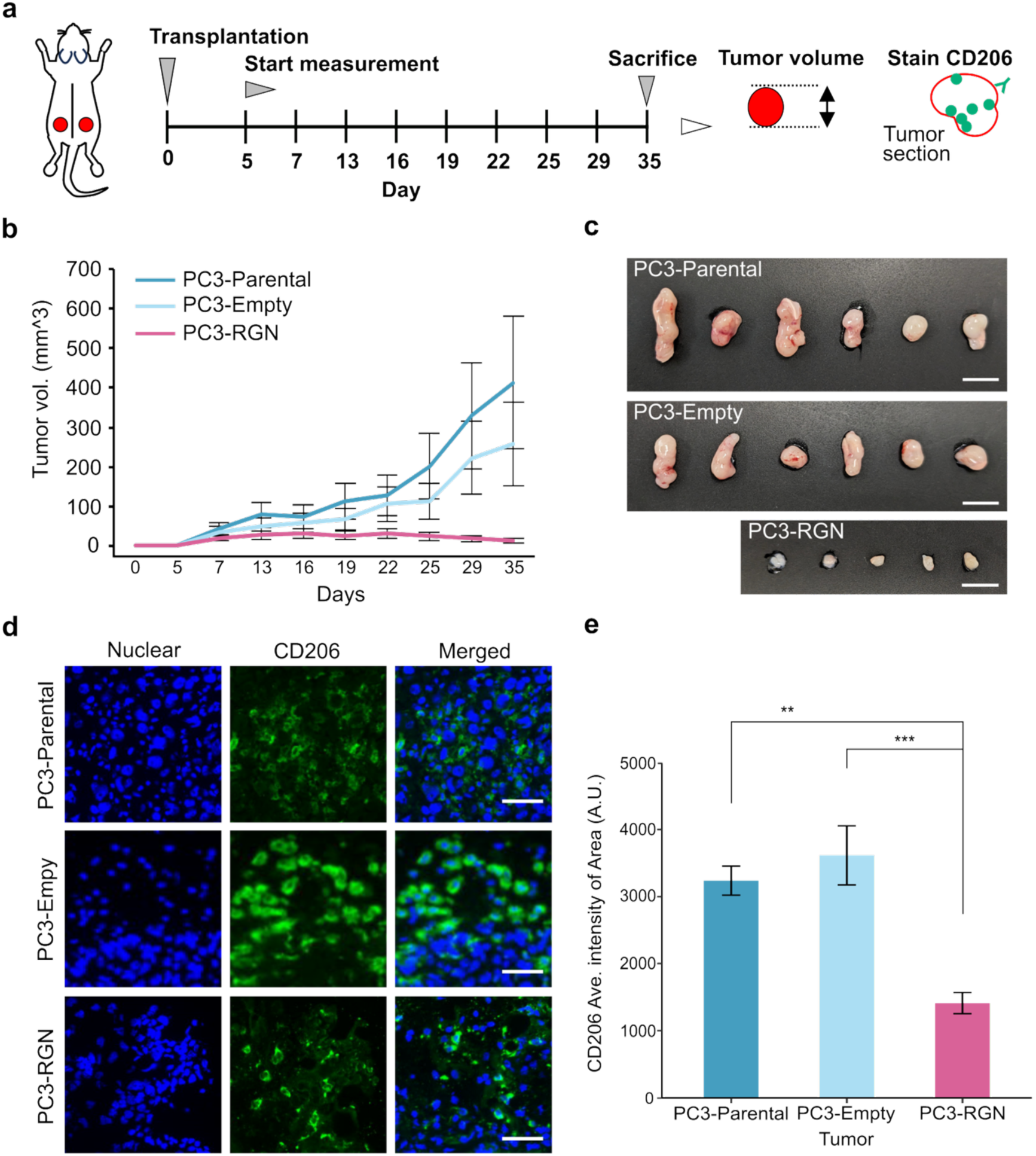
CD206 macrophage suppression in PC3-RGN tumors. **a.** *In vivo* tumor formation protocol. Cells were transplanted two sites per mouse. (3 mice for each group) **b.** Tumor volume over 35 days. PC3-Parental and PC3-Empty: n = 6; PC3-RGN: n = 5 (one tumor failed to develop). **c.** Photographs of excised tumors on day 35 for each group. Scale bar: 10 mm. **d.** Immunofluorescence staining of CD206+ macrophages in PC3-Parental (top row), PC3-Empty (middle row), and PC3-RGN (bottom row) tumors. Left: DAPI staining (nuclei, blue); Middle: anti-CD206 staining (green); Right: merged image. Scale bar: 50μm. **e.** Quantification of CD206+ macrophage intensity normalized to tumor area across all three groups. (PC3-Parental, n = 6; PC3-Empty, n = 6; PC3-RGN, n = 5). Statistical analysis: one-way ANOVA followed by the Tukey–Kramer post-hoc test. PC3-RGN vs PC3-Parental: ***P < 0.001; PC3-RGN vs PC3-Empty: **P < 0.01.

### RGN suppresses proliferation of PC3 cancer cells

To elucidate the effects of RGN in cells, an RGN-expressing PC3 cell line was generated by transfecting empty vector or RGN expression constructs into parental PC3 cells. Clonal cell lines were derived from empty vector-transfected cells (#1–#4) and RGN expression construct-transfected cells (#1–#3). RGN expression in all sublines was confirmed by western blot analysis (Supplementary Fig. 1a), with RGN levels quantified using β-actin (Supplementary Fig. 1b) or GAPDH (Supplementary Fig. 1c) as loading controls. Among the empty vector-transfected sublines, subline #3 (designated PC3-Emp) exhibited low RGN expression comparable to parental PC3 cells and was selected for subsequent experiments. Among the RGN expression construct-transfected sublines, subline #3 (designated PC3-RGN) exhibited the highest RGN expression compared to sublines #1 and #2 and was selected for further analysis. Cell proliferation assays over 5 days revealed that PC3-RGN cells demonstrated a significant reduction in proliferation compared to PC3-Emp cells (0.6-fold reduction, p < 0.01, Student’s t-test; Supplementary Fig. 1d). In contrast, the proliferation rate of PC3-Emp cells was not significantly different from that of parental PC3 cells. These findings indicate that RGN expression suppresses PC3 cancer cell proliferation.

### CD206⁺ M2-like macrophages are reduced in RGN-expressing PC3 tumors

To investigate tumorigenesis in established cell lines, we conducted *in vivo* analyses. Parental PC3, PC3-Empty, or PC3-RGN cells were subcutaneously transplanted into two dorsal sites per mouse (Fig. 2a). Tumor volumes were monitored for a 35-day period (Fig. 2b). All tumors were excised on day 35; however, one of the six PC3-RGN transplantation sites failed to develop a tumor (Fig. 2c). The mean PC3-RGN tumor volume was 13 mm3 (± 5.3), while the mean parental PC3 and PC3-Empty tumor volumes were 413.1 mm3 (± 168.7) and 258.2 mm3 (± 105.4), respectively. On day 35, tumors derived from PC3-RGN cells were significantly smaller than those in the other groups (Supplementary Fig. 2a). The tumors were subjected to immunofluorescence with anti-human vimentin (Supplementary Fig. 2b). The PC3 tumor cells were clearly demarcated from the surrounding murine cells, with murine cell infiltration confined to localized regions. Notably, PC3-RGN tumors were significantly smaller than those from parental PC3 and PC3-Empty cells (Supplementary Fig. 2a), an effect that exceeded the expected reduction based on the in vitro proliferation defect alone (Supplementary Fig. 1d). We postulated that cell-cell communication between RGN-expressing cells and TME modulates tumorigenesis.

We subsequently focused our investigation on tumor-associated macrophages, given their pivotal role as key regulators of tumorigenesis. Harvested tumor tissues were subjected to immunofluorescence staining with an anti-CD206 antibody, a marker for M2 macrophage polarization (Fig. 2d). The CD206 fluorescence intensity normalized to the tumor area within PC3-RGN tumors was significantly reduced compared to those observed in parental PC3 or PC3-Empty tumors (Fig. 2e). Additionally, harvested tumor tissues were stained with an anti-CD80 antibody, a marker for M1 macrophage polarization (Supplementary Fig. 2c); CD80 signals were below the detection limit under the present conditions, and thus the M1 status could not be reliably evaluated. The PC3-RGN subline exhibited a 0.6-fold reduction in proliferation compared to the other cell lines *in vitro* (Supplementary Fig. 1d). These results indicate that RGN expression in PC3 cells was associated with reduced cell proliferation *in vitro* and with a reduced CD206⁺ M2-like macrophage signal in the resulting tumors.

### RGN is contained in EVs derived from PC3-RGN cells

We focused our investigation on EVs derived from the PC3-RGN subline as potential mediators of tumor suppression. EVs were purified and concentrated from the conditioned media of each cell line using differential ultracentrifugation, according to established protocols ^10^. Intensity-weighted size distributions were obtained by dynamic light scattering with the ELSZneo instrument. DLS provides an intensity-weighted average and assumes particle uniformity; the values below are therefore used only to compare the three preparations with each other and not as absolute particle sizes or concentrations. The modal hydrodynamic diameters of EVs from parental PC3, PC3-Empty, and PC3-RGN sublines were 165.9 nm, 161.5 nm, and 167.6 nm, respectively (Fig. 3a), indicating similar size distributions across the three cell lines. The morphology of the EVs was visualized using transmission electron microscopy with negative staining (Fig. 3b), revealing a characteristic cup-shaped morphology consistent with vesicle structures ^24^. Western blot analysis was performed on cell lysates and purified EVs (Fig. 3c). RGN was detected exclusively in EVs derived from the PC3-RGN subline, and not in those from the parental PC3 or PC3-Empty sublines. In accordance with MISEV2023, CD63 was used as a transmembrane EV marker (Category 1), β-actin and GAPDH as cytosolic proteins (Category 2), and cytochrome C as a marker of intracellular compartments other than the plasma membrane and endosomes (Category 4). CD63 exhibited a robust signal in EVs compared to cell lysates, particularly in EVs from the PC3-RGN subline. β-actin and GAPDH were below the limit of detection in EVs derived from the PC3-RGN. Cytochrome-C was not detected in any of the EV samples, indicating a low level of mitochondrial co-isolation. These findings indicate that RGN expression in cells is accompanied by a change in the protein composition of the secreted EVs.

**Figure 3.**
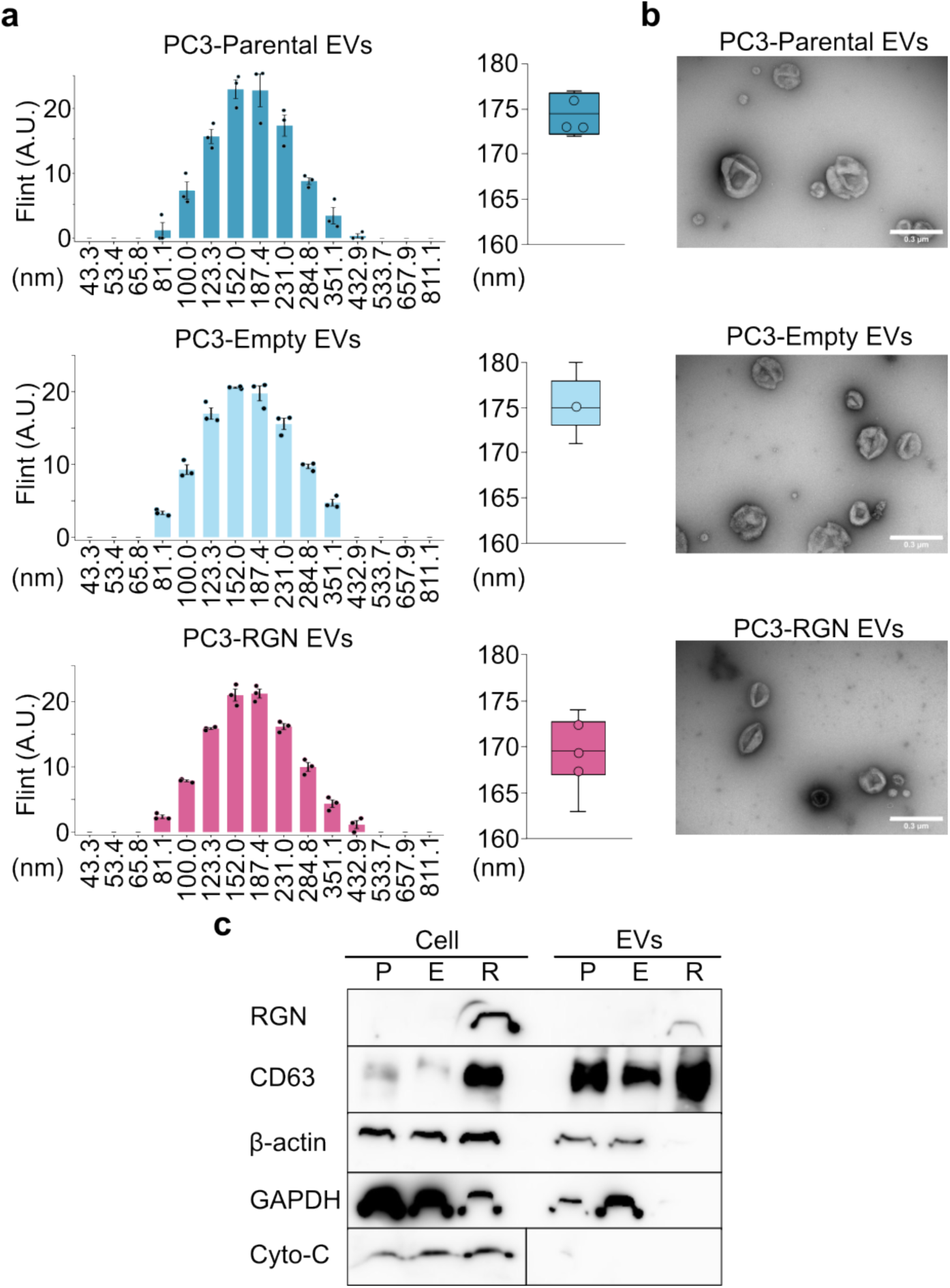
Characterization of extracellular vesicles (EVs) isolated from sub-cell lines. **a.** Size distribution analysis of representative EVs presented as a histogram, and size distribution of EVs from three independent EV preparations presented as a box plot. (n = 3 for each group) **b.** Transmission electron microscopy image of isolated EVs. Scale bar: 300 nm. **c.** Western blot analysis of RGN, CD63, β-actin, GAPDH, and Cytochrome-C protein expression in sub-cell line lysates and isolated EVs, shown as a representative blot. Error bars represent standard deviation (S.D.).

### RGN-containing EVs suppress M2 macrophage polarization

To investigate the role of EVs in suppressing M2 polarization, J744α1 cells, a mouse macrophage cell line, were treated with 1 μg/mL of EVs derived from parental PC3, PC3-Empty, or PC3-RGN cells for two days. iNOS (M1 marker) and Arg-1 (M2 marker) expression in J744α1 cells were measured using real-time PCR (Fig. 4a and b). Arg-1 expression was suppressed in cells treated with PC3-RGN EVs, but increased in cells treated with parental PC3 or PC3-Empty EVs. iNOS expression tended to increase in EV-treated cells compared to untreated cells, although not significantly. To elucidate the signaling pathway involved in M2 polarization, western blotting was performed (Fig. 4c). J744α1 cells were treated with IFN-γ (M1 polarization) or IL-4 (M2 polarization). IFN-γ activated p-STAT1 (Y701), p-STAT1 (S727), and p-ERK1/2, indicating M1 polarization. IL-4 suppressed p-AKT1, and p-ERK1/2, indicating M2 polarization. PC3-RGN EVs reduced the p-AKT1/AKT1 and p-ERK1/2/ERK1/2 ratios († in Fig. 4c). Akt1, a component of the PI3K pathway, is a key factor in M2 polarization ^25^. These results suggest that the reduction in M2 marker expression by PC3-RGN EVs is accompanied by decreased AKT1 and ERK1/2 phosphorylation.

**Figure 4.**
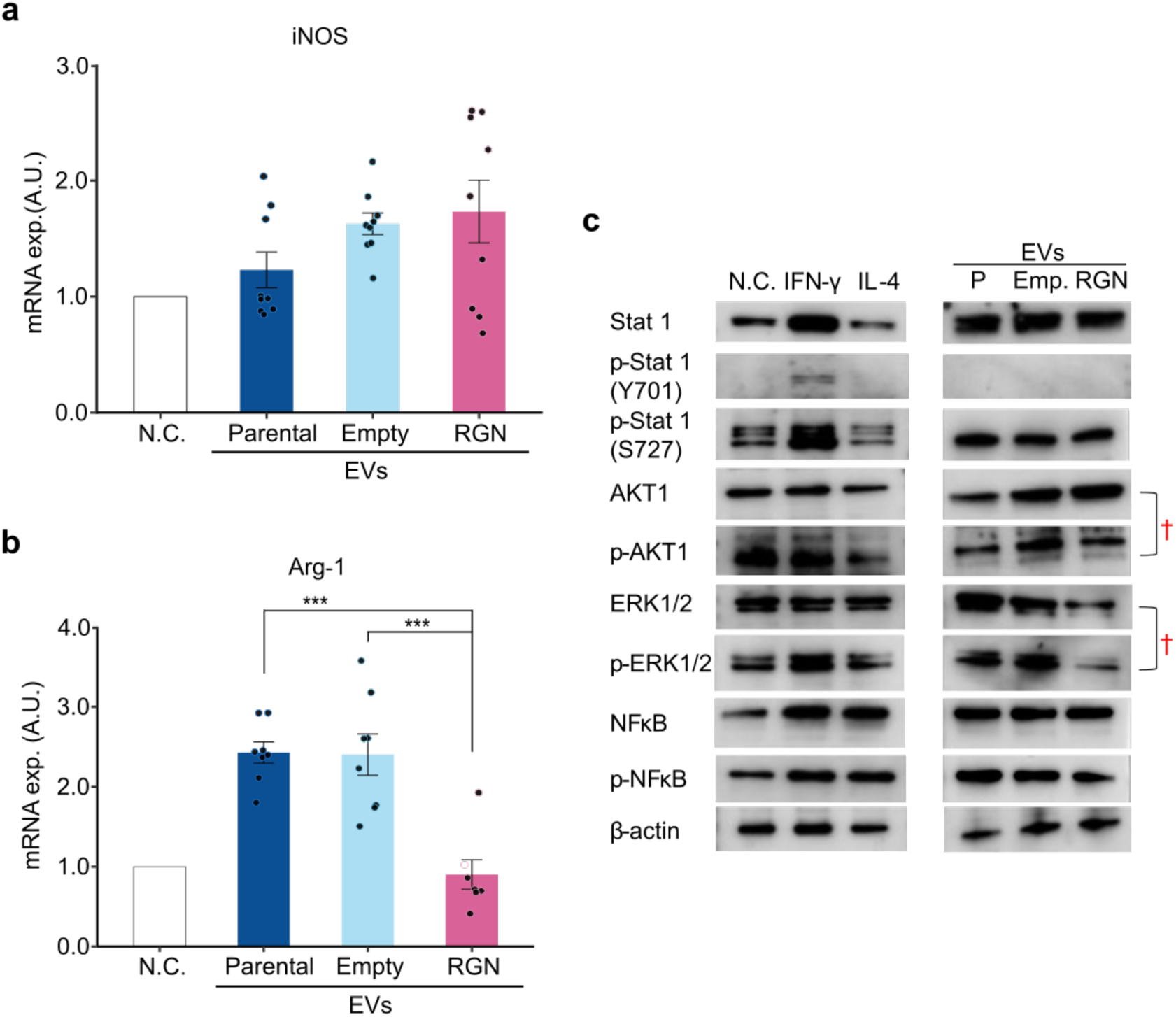
PC3-RGN-derived EVs decrease Arg-1 expression in J744α1 cells. **a-b.** iNOS (A) (n = 9 for each group) and Arg-1 (B) (N.C., n = 9, Parental, n = 8; Empty, n = 8; RGN, n = 7) mRNA levels after treatment with EVs. Data are from three independent experiments, each performed in triplicate. Statistical analysis: one-way ANOVA followed by the Tukey–Kramer post-hoc test. **c.** Western blot showing Stat1, p-Stat1 (Y701), p-Stat1 (S727), AKT1, p-AKT1, ERK1/2, p-ERK1/2, NFκB, p-NFκB, and β-actin in J744α1 cell lysates. Left: untreated (NC), IFN-γ, and IL-4 treated cells. Right: EV-treated cells. P: parental PC3 cells. †: indicates proteins with altered expression levels in this representative blot. \*\*\**P* < 0.001. Error bars represent standard deviation (S.D.).

### RGN suppresses M2 polarization via signaling pathway modulation

PC3-RGN EVs suppressed M2 polarization by blocking AKT1 and ERK signaling. To identify the key factor in M2 polarization suppression, we focused on the RGN protein. J744α1 cells were transfected with recombinant RGN. Immunofluorescence was performed on transfected cells using an anti-RGN antibody (Fig. 5a). Recombinant RGN was taken up by J744α1 cells (Fig. 5b). After two days, western blotting was performed on RGN-transfected J744α1 cell proteins (Fig. 5c). Recombinant RGN suppressed p-AKT1/AKT1 and p-ERK1/2/ERK1/2 († in Fig. 5c). These results suggest that RGN itself is sufficient to reproduce the changes in AKT1 and ERK1/2 phosphorylation observed with RGN-containing EVs. Recombinant RGN transfection yielded similar results to RGN-containing EV treatment.

**Figure 5.**
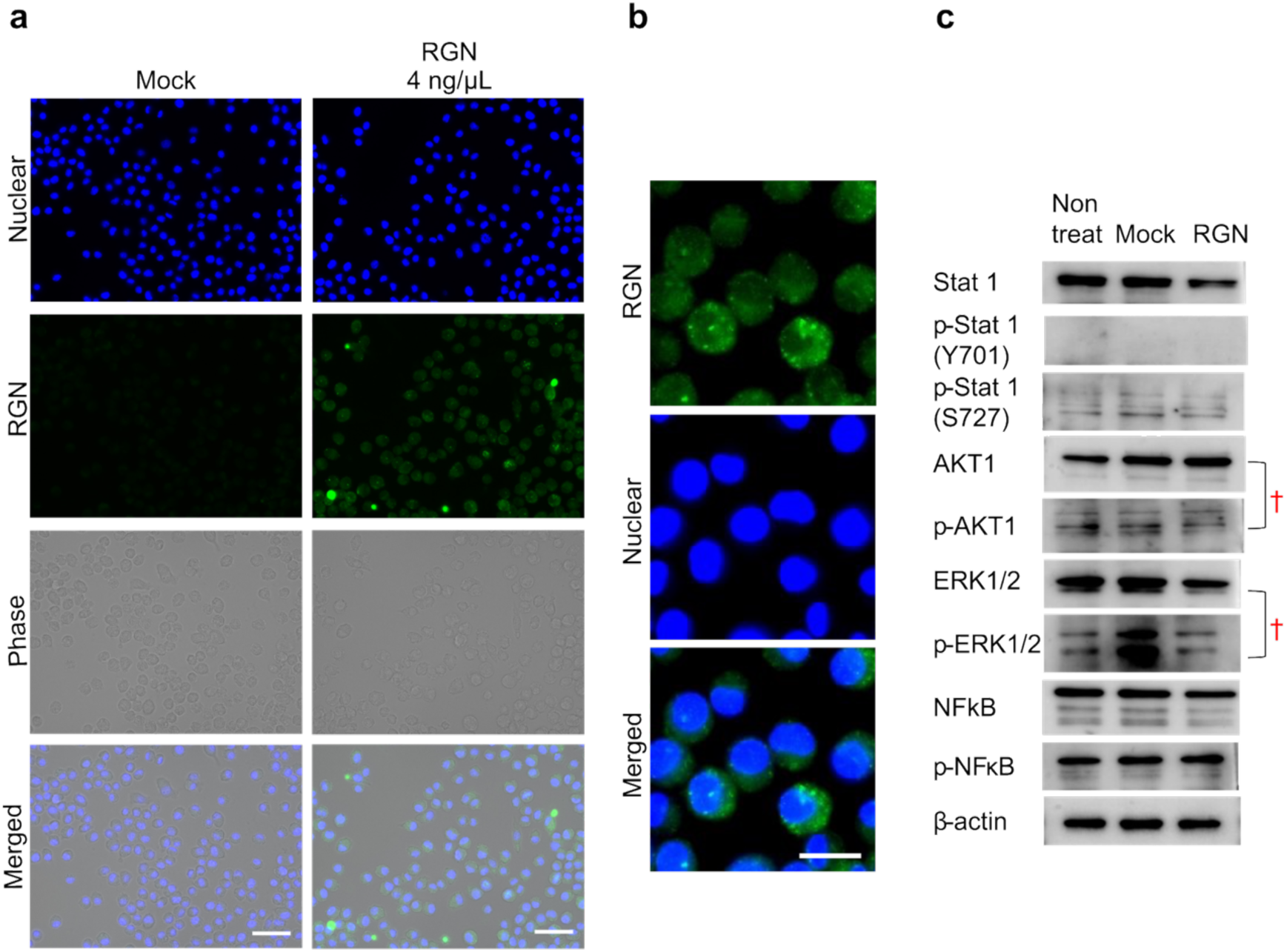
RGN suppresses p-AKT and p-ERK. **a.** RGN immunofluorescence (J744α1; recombinant RGN transfection). Mock: transfection reagent only. Scale bar: 50 μm. **b.** Enlarged RGN immunofluorescence (J744α1; recombinant RGN transfection). Scale bar: 20 μm. **c.** Western blot: Stat1, p-Stat1 (Y701), p-Stat1 (S727), AKT1, p-AKT1, ERK1/2, p-ERK1/2, NFκB, p-NFκB, β-actin (J744α1 lysate), shown as a representative blot. †: indicates proteins with altered expression levels in this representative blot.

**Figure 6.**
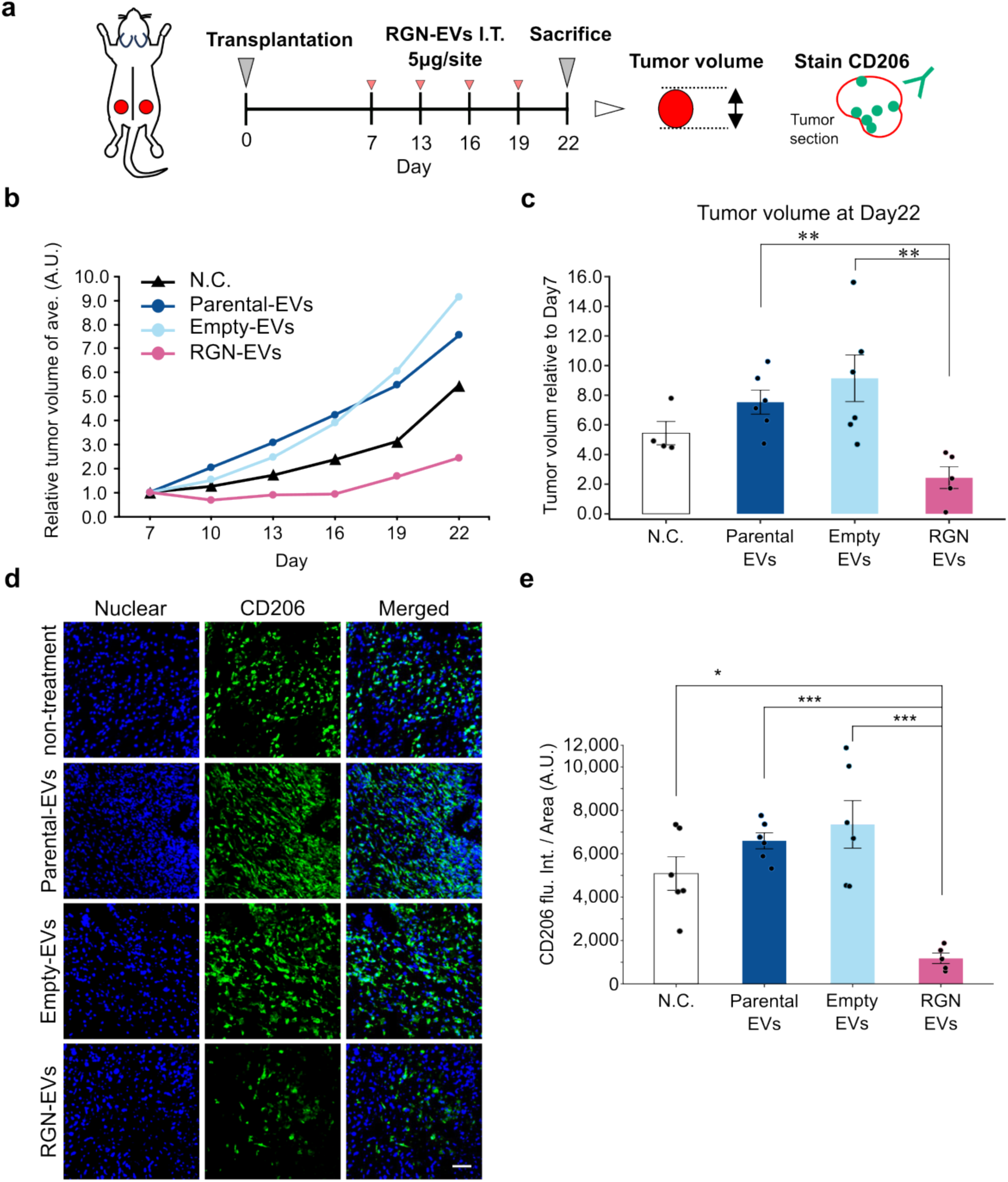
Antitumor effects of PC3-RGN EVs. **a.** *In vivo* EV treatment protocol. Cells were transplanted two sites per mouse. (3 mice for each group) EVs were injected intratumorally at 5 μg per site on days 7, 13, 16, and 19. **b.** Tumor volume over 22 days, expressed relative to day 7. (6 sites for each group) **c.** Tumor volume at day 22, expressed relative to day 7 (arbitrary units). (6 sites for each group) **d.** CD206 immunofluorescence (scale bar: 50 μm). **e.** CD206 intensity normalized to the tumor area. (N.C., n = 6; Parental-EVs, n = 6; Empty-EVs, n = 6; RGN-EVs, n = 5) Statistical analysis for c and e: one-way ANOVA followed by the Tukey–Kramer post-hoc test. \**P* < 0.05, \*\**P* < 0.01, \*\*\**P* < 0.001. Error bars represent standard deviation (S.D.).

### PC3-RGN EVs suppress tumor growth and reduce CD206^+^ M2 -like macrophages *in vivo*

Encouraged by *in vitro* results, we investigated the antitumor effects of PC3-RGN EVs, containing RGN, in PC3 tumor-bearing BALB/c-nu/nu mice. Parental PC3 cells were transplanted subcutaneously at two sites per mouse. Tumor volume was observed for 22 days (Fig. 6a). EVs were administered into the tumor approximately every seven days at 5 μg/site for four injections (Fig. 6b). PC3-RGN EV-treated tumors were significantly smaller than parental PC3 or PC3-Empty EV-treated tumors on day 22 (Fig. 6c). To confirm M2 macrophage polarization, we performed CD206 immunofluorescence (Fig. 6d). CD206-positive cell fluorescence in PC3-RGN EV-treated tumors was significantly lower than in parental PC3 or PC3-Empty EV-treated tumors (Fig. 6e). PC3-RGN EV treatment suppressed tumor growth and was associated with a reduced CD206⁺ M2-like macrophage signal, suggesting that M2 suppression may contribute to the antitumor effect.

## Discussion

In the current study, we found that regucalcin-containing EVs are associated with reduced tumor growth and reduced M2 macrophage polarization in TME (Fig. 7). Moreover, the downregulation of regucalcin in malignant tissues is correlated with advanced tumor stages and reduced patient survival. We also showed that regucalcin overexpression suppressed PC3 cell proliferation *in vitro* and EVs derived from PC3-RGN cells contained RGN. Interestingly, RGN-containing EVs suppressed M2-polarized tumor-associated macrophages, suggesting that RGN is sufficient to reproduce this effect on macrophages. In vivo, PC3-RGN EV treatment suppressed tumor growth and was associated with reduced CD206⁺ M2-like macrophage infiltration. Together with the in vitro findings, these results suggest that RGN-containing EVs suppress tumor growth, at least in part, by attenuating M2 macrophage polarization. Additionally, *in vitro* experiments showed that the attenuation of M2 macrophage polarization by RGN-containing EVs was accompanied by decreased AKT1 and ERK1/2 phosphorylation.

**Figure 7.**
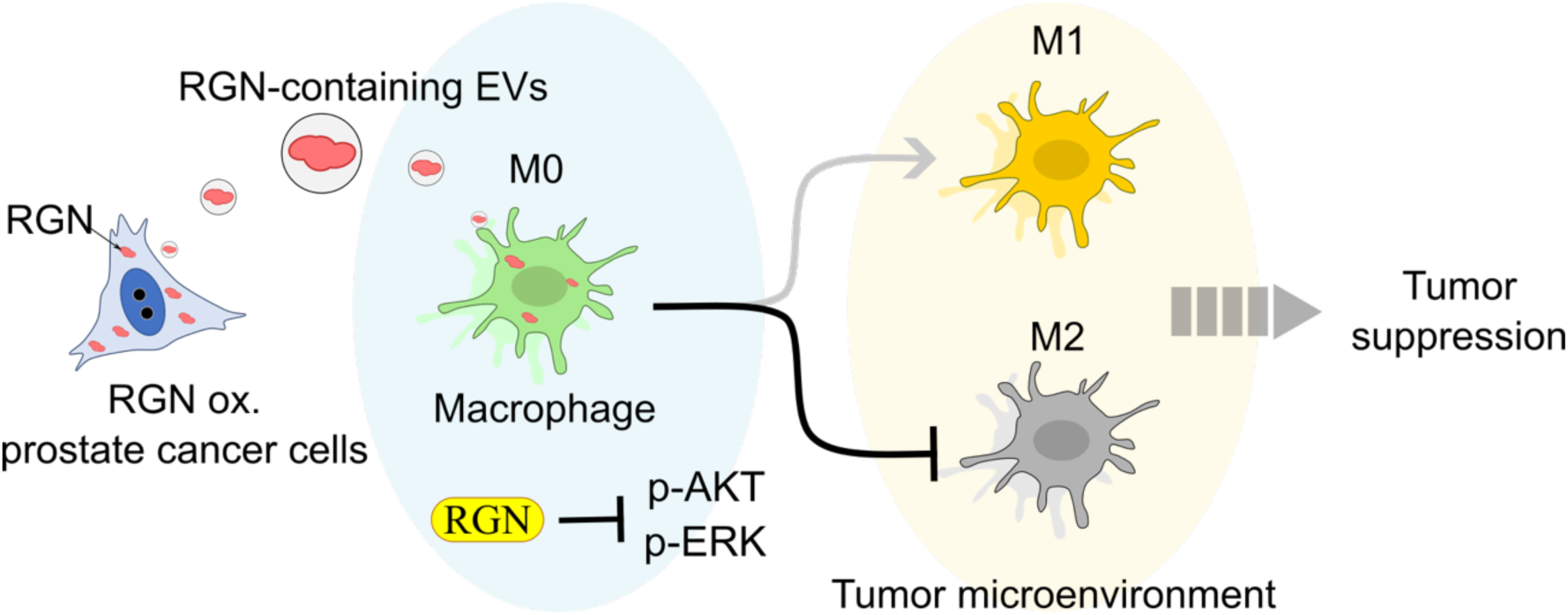
RGN-containing EVs uptake by macrophages leads to the suppression of M2 polarization through signaling pathways.

A key finding of our research is the identification of RGN within EVs, which, to the best of our knowledge, is the first report. We observed significant attenuation of CD206-expressing M2 macrophages in RGN-expressing PC3 tumors. M2 macrophages, characterized by elevated CD206 expression ^26^, are established drivers of tumor progression through proangiogenic ^27^ and immunosuppressive mechanisms ^28,29^. Macrophage polarization toward the M2 phenotype, encompassing CD206 upregulation, is frequently mediated by transforming growth factor-beta (TGF-β) ^30–33^, a pleiotropic cytokine that modulates diverse cellular processes, including immune cell differentiation. TGF-β signaling is initiated by ligand (TGF-β1, TGF-β2, and TGF-β3) binding to a heteromeric receptor complex consisting of type II (TGFBR2) and type I (TGFBR1/ALK5) serine/threonine kinase receptors ^31^. This receptor complex subsequently activates intracellular signaling cascades, primarily through the phosphorylation of SMAD2 and SMAD3, which then assemble with SMAD4 and translocate to the nucleus ^34–36^. This SMAD complex, along with other transcription factors, regulates target gene expression, including CD206 ^34,37^. TGF-β can also activate MAPK and PI3K/AKT pathways, contributing to CD206 expression ^37^. Consistent with these pathways, RGN diminishes Ras, PI3K, Akt, and MAPK levels, which are key signaling molecules in PC-3 cell growth ^9^. As illustrated in Figure 7, our findings suggest that RGN-containing EVs inhibit M2 polarization, at least partially, by interfering with these signaling pathways.

The observation that RGN-containing EVs are associated with a change in macrophage phenotype supports further evaluation of an EV-based approach ^38^. Direct delivery of RGN to macrophages in vivo was not demonstrated in this study. The established inverse correlation between RGN expression and malignant progression reinforces the rationale for utilizing RGN-containing EVs to restore RGN function and impede tumor growth. Critically, the suppression of M2 macrophage polarization by RGN-containing EVs is highly significant, given the well-documented role of M2 macrophages in promoting tumor progression and establishing an immunosuppressive milieu. Mechanistic insights into EV-mediated macrophage polarization modulation provide a valuable understanding of tumor-immune system interactions. This discovery aligns with established paradigms demonstrating macrophage polarization modulation as a promising cancer therapeutic strategy ^39^. Several limitations should be noted. Our in vivo data are correlative and do not establish that M2 suppression causally drives the reduced tumor growth. Although recombinant RGN reproduced the EV effect on macrophages—supporting that RGN is sufficient—its necessity within EVs was not tested. The magnitude of the in vivo effect also exceeded what the in vitro proliferation data would predict: PC3-RGN cells showed a 0.6-fold reduction in proliferation relative to control cells, whereas the mean PC3-RGN tumor volume on day 35 was approximately 30-fold smaller than that of parental PC3 tumors. Reduced intrinsic proliferation therefore does not account for the in vivo phenotype on its own, and host-dependent processes that were not examined here—including altered angiogenesis, recruitment of other innate immune populations, or differences in initial tumor establishment—may also contribute. A further limitation concerns the assessment of macrophage phenotype: CD80 staining was below the detection limit in our sections, so we could not determine whether the reduction in CD206⁺ cells reflects a shift in the M1/M2 balance or a decrease in overall macrophage infiltration. In addition, the EVs were prostate cancer–derived and based on a single cell line in a cross-species setting, so cargo safety and generalizability require further study. The in vivo experiments used BALB/c nu/nu mice, which lack functional T cells; any contribution of T cell–mediated antitumor immunity could not be assessed, and validation in immunocompetent or syngeneic models will be required. In addition, the RGN content of the EV preparations was not quantified in absolute terms, so the concentration of recombinant RGN used for transfection (4 ng/μL) cannot be directly related to the amount of RGN delivered by EVs; the recombinant protein experiment should therefore be regarded as a test of whether RGN is sufficient to produce the observed signaling changes, rather than as a dose-matched comparison.

Though the effects of RGN-containing EVs on other immune cells within the TME warrant further study ^40–42^, this study focused on macrophages. Future research could explore interactions with T cells, dendritic cells, and natural killer cells, potentially revealing additional antitumor mechanisms and synergistic effects with immunotherapies. We acknowledge our study does not demonstrate EV-mediated targeted delivery with absolute specificity. While EVs offer theoretical potential for enhancing treatment specificity and minimizing off-target effects, achieving this remains a significant challenge ^43^.

In summary, our findings suggest that RGN-containing EVs represent a candidate strategy for cancer treatment that warrants further evaluation. By harnessing EV properties and RGN’s anti-tumorigenic activity, we have identified a candidate pathway for EV-based cancer therapy warranting further investigation. Continued research, including causal validation by macrophage depletion or CD206 knockdown and evaluation in immunocompetent models, will be required to determine whether EV-based treatments of this type can improve patient outcomes.

## Acknowledgements

We express our sincere gratitude to the study participants for their generous donation of time. We thank Ms. Saki Horie for her technical assistance. We thank Dr. Tomiyasu Murata, Faculty of Pharmacy at Meijo University, for the kind gift of the RGN expression vector. We also thank the Science Research Center of Yamaguchi University for Institute of Gene Research, Institute for Biomedical Research and Institute of Life Science and Medicine. This work was the result of using research equipment shared in MEXT Project for promoting public utilization of advanced research infrastructure (Program for supporting construction of core facilities), Grant Number JPMXS0440400024.

## Fundings

This work was supported in part by a Grant-in-Aid for Research Activity Start-up (No. 21K21219), Early-Career Scientists (No.22K17826) from Japan Society of the Promotion of Science (JSPS). This work was supported by Japan Science and Technology Agency (JST), ACT-X Grant Number JPMJAX222B, Japan. This work was supported in part by a Grant-in-Aid for Yamaguchi University Fund, Start-up Research Fund, New frontier project-2021, and FOCS 2023 (Yamaguchi University Original-University Fund). This work was supported in part by a Grant-in-Aid for the Ube City Next-Generation Researchers Project. The funding bodies were not involved in the design of the study and collection, analysis, and interpretation of data and in writing the manuscript.

## Author contribution

K.T. and N.T. conceived and designed this study. K.T., N.T., and R.O. contributed to figure preparation and data presentation. R.O., and N.T. performed the animal experiments. R.O., K.T., and N.T. performed data analysis and interpretation. T.Y. and M.Y. provided helpful discussions. All authors reviewed and edited the manuscript. The manuscript was finalized by N.T. with the assistance of all authors, and all authors approved the final manuscript.

## Disclosure of interest

The author N.T. is the founder and director of ADDVEMO, Inc. The author N.T. receives research found from ADDVEMO Inc. These funder had no role in this study design, data collection, analysis, interpretation of data, the writing of the report, or the decision to submit the article for publication. The authors R.O., K.T., T.Y., and M.Y. declare no conflict of interest.

## Notes

### Competing Interest Statement

The author N.T. is the founder and director of ADDVEMO, Inc. The author N.T. receives research found from ADDVEMO Inc. and MIZUTA Seisakusho, Inc.. These funders had no role in this study design, data collection, analysis, interpretation of data, the writing of the report, or the decision to submit the article for publication. The authors R.O., K.T., T.Y., and M.Y. declare no conflict of interest.

