## Supplementary Figure for "Regucalcin-containing extracellular vesicles suppress M2 macrophage polarization and attenuate tumor progression in vivo"

Naoomi Tominaga, Ph.D.

Department of Clinical Laboratory Science, Faculty of Health Sciences, Yamaguchi University, Graduate School of Medicine, 1-1-1 Minami-kogushi, Ube, Yamaguchi, 755-8505, Japan

### Keywords

Extracellular vesicles, Regucalcin, Macrophage, Tumor associated macrophage, anti-tumor therapy

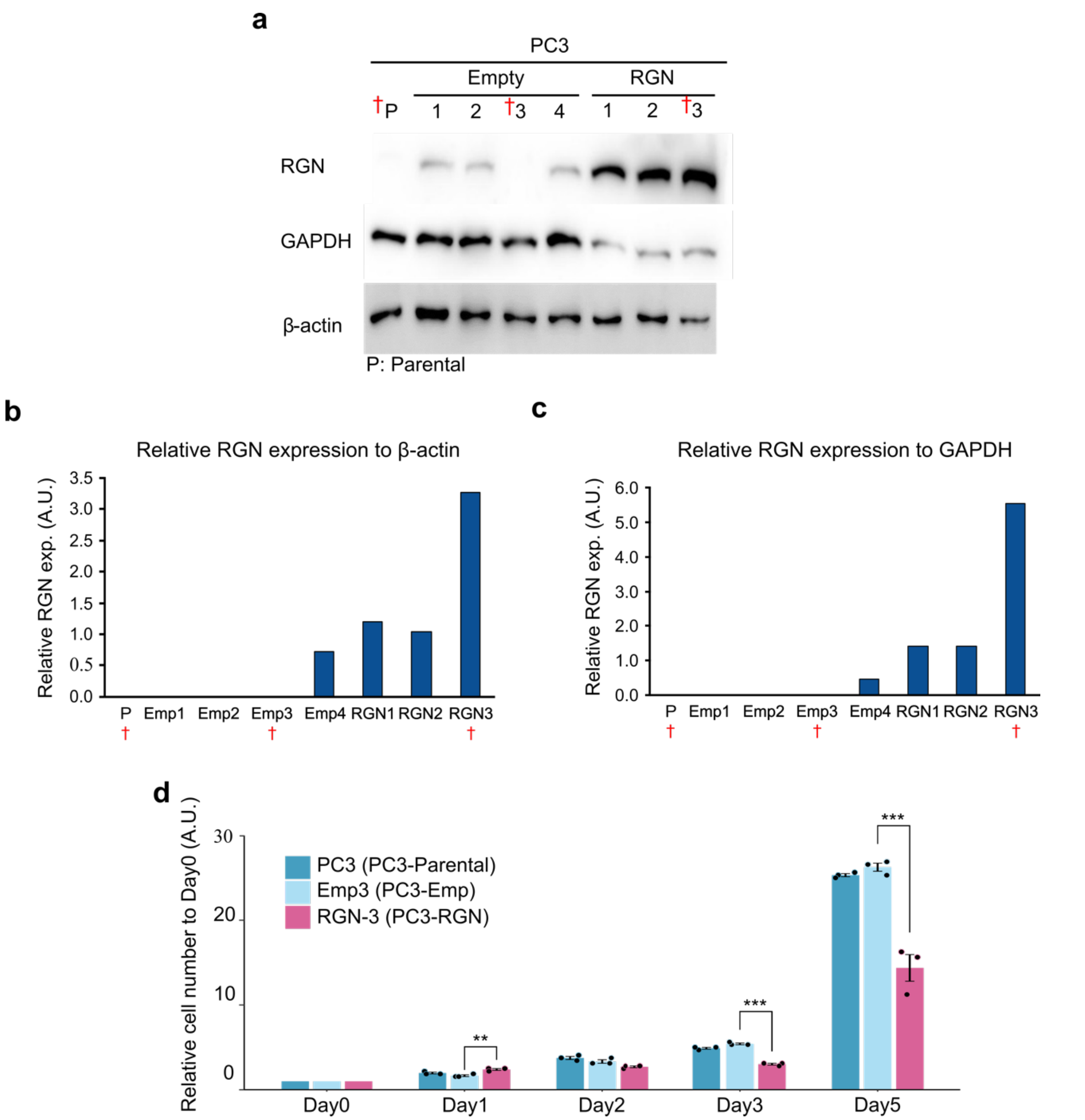


**Supplementary Figure 1.**

a. Western blot analysis of RGN, GAPDH, and β-actin expression in parental PC3 cells (P), empty vector-transfected sub-cell lines (#1-#4), and RGN expression construct-transfected sub-cell lines (#1-#3). GAPDH and β-actin served as loading controls. † indicates cell lines used. **b.** Quantification of RGN expression intensity normalized to β-actin (loading control) across all sub-cell lines. † indicates selected cell lines. **c.** Quantification of RGN expression intensity normalized to GAPDH (loading control) across all sub-cell lines. † indicates selected cell lines. **d.** Cell proliferation assay of PC3-Parental, PC3-Emp (#3), and PC3-RGN (#3) measured over 5 days, expressed relative to day 0. (n = 6 per group; Student's t-test, $^{**}P < 0.01$). † indicates selected cell lines. Error bars represent standard deviation (S.D.).

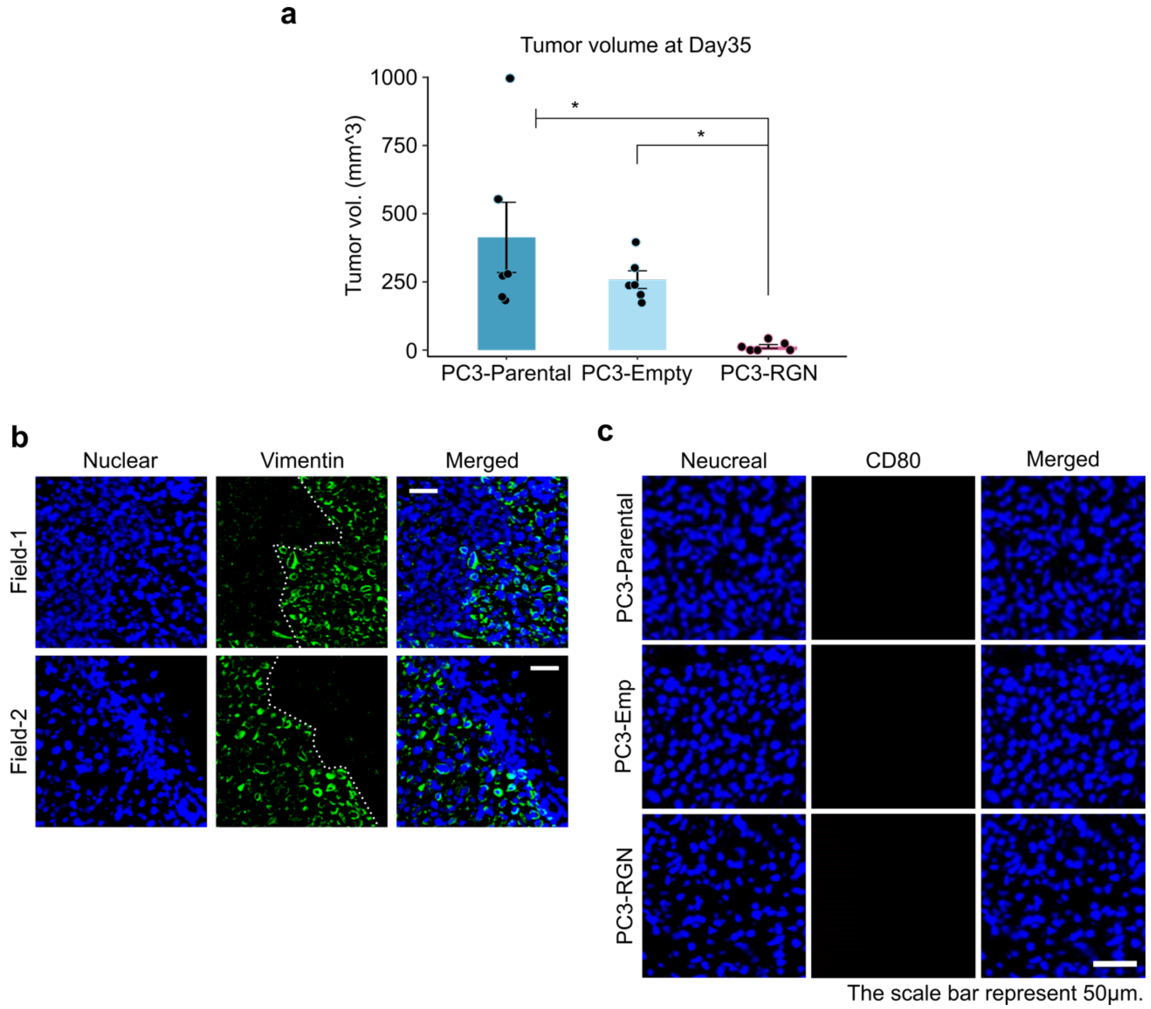


**Supplementary Figure 2.**

**a.** Tumor volume at day 35. (n = 6 per group; one implantation site in the RGN group did not develop a tumor and was included with a tumor volume of 0 mm³) Statistical analysis: one-way ANOVA followed by the Tukey–Kramer post-hoc test. **b.** Vimentin immunofluorescence in parental PC3 tumor (scale bar: 50 μm). Green denotes vimentin, while white dotted marks indicate the boundary. **c.** Immunofluorescence image of tumor stained with anti-CD80. Scale bar represents 50 μm. *$P < 0.05$, Error bars represent standard deviation (SD).
